# Improving peptide vaccine manufacturability without sacrificing immunogenicity: substitution of methionine and cysteine with oxidation-resistant isosteres

**DOI:** 10.64898/2026.01.20.699901

**Authors:** Andrew S. Ishizuka, Christopher M. Garliss, Robert N. Goddu, Maria Merolle, Alec Schrager, Andrei Ramirez-Valdez, Qiuyin Ren, Faezzah Baharom, Matthew Essandoh, Nicholas G. Palacorolla, John P. Finnigan, Daniel C. Douek, Nina Bhardwaj, Robert A. Seder, Geoffrey M. Lynn, David R. Wilson

**Author notes:** Corresponding authors: David R. Wilson and Geoffrey M. Lynn.

## Abstract

Vaccines comprising peptide antigens for inducing T cell immunity are being developed for a broad range of therapeutic applications including prevention and treatment of cancer, autoimmunity, and infectious diseases. However, many peptide antigens contain cysteine and/or methionine, which are prone to form oxidation products that can present challenges to manufacturing and reduce biological activity. To address this challenge, we introduced oxidation resistant (OXR) antigens wherein the cysteine and methionine residues of naturally occurring, wild type (WT) peptide antigens are substituted with isosteric residues that are structurally related but omit the oxidation-prone sulfur atom. Our results showed that vaccination with OXR antigens substituting cysteine and methionine with isosteres alpha-aminobutyric acid and norleucine, respectively, induced immune responses to the WT antigen that were equivalent or higher than those induced by vaccination with WT antigens. T cell responses were not affected by the position of the amino acid substitutions indicating that the isosteres do not negatively impact major histocompatibility complex (MHC) binding or T cell recognition. The T cells induced were high quality and associated with anti-tumor efficacy *in vivo*. Interestingly, substitution of cysteine with serine, which replaces the sulfur for an oxygen, did not yield cross-reactive T cell responses, highlighting the high degree of molecular discernment of peptide-MHC processing and presentation. In sum, OXR antigens provide a generalizable strategy for eliminating sulfur oxidation products and improving the manufacturability and shelf-life of peptide-based vaccines without affecting desired biologic activity.

## Introduction

Vaccines comprising synthetic peptide antigens are currently under development for a variety of medical applications, including prevention or treatment of infectious diseases, cancer, and autoimmunity^1^. Most peptide-based vaccine approaches rely on peptide antigens based on naturally occurring proteins from pathogens or the human proteome that are produced synthetically as shorter fragments typically between 8-35 amino acids in length^2^. Each peptide antigen includes a short sequence, referred to as an epitope, which binds major histocompatibility complex (MHC) molecules and is recognized by T cells as a peptide-MHC complex (**Fig. 1a**)^3,4^. Amino acid residues within a given epitope interact either with the T cell receptor or MHC (as anchor residues), and thus, the identity and structure of each residue is critical to proper recognition of epitopes by T cells^5^.

**Fig. 1:**
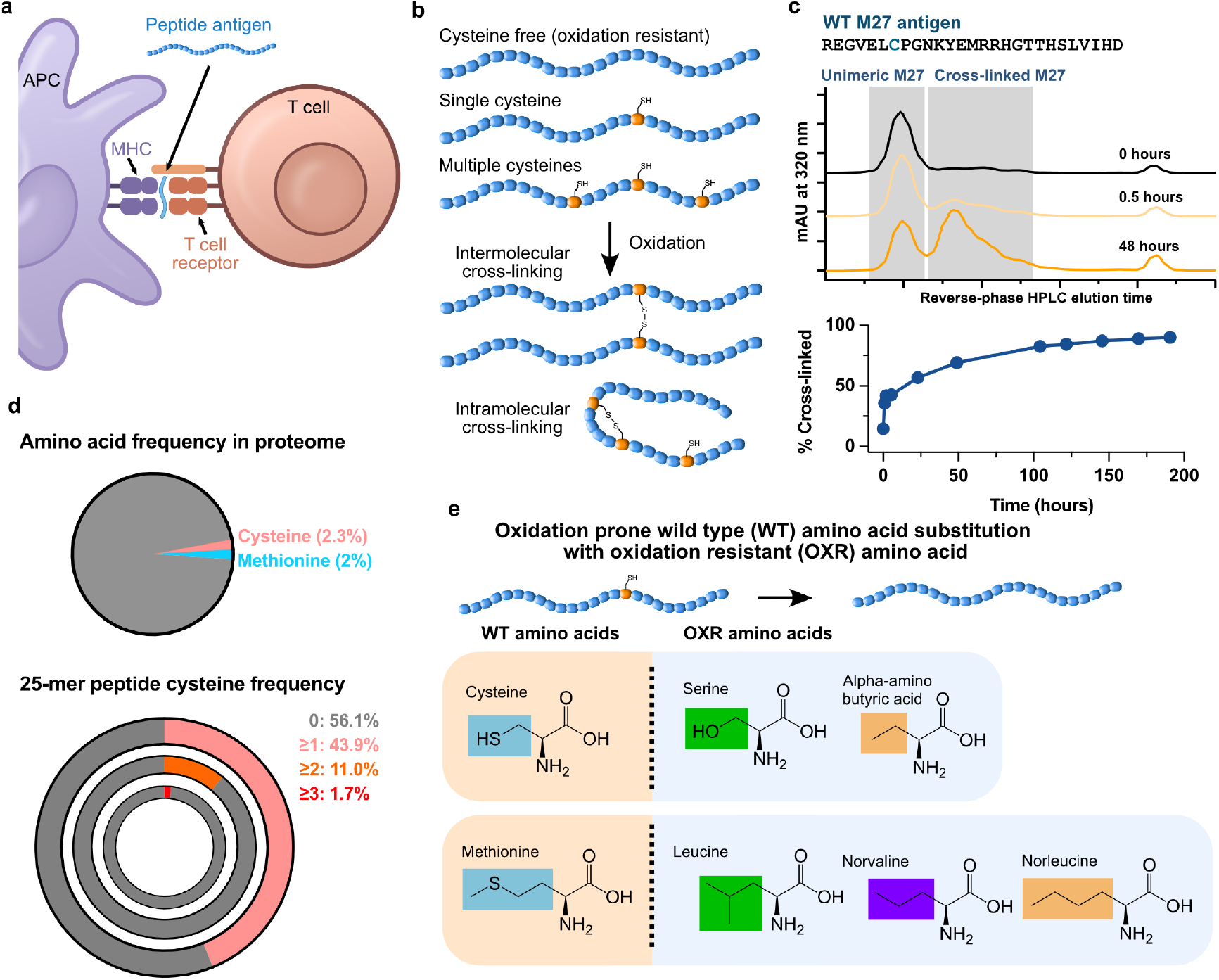
Oxidation-prone amino acids in peptide antigens. **(a)** Cartoon schematic depicting T cell receptor recognition of cognate peptide-MHC complex as presented by an APC. **(b)** Cartoon schematic depicting how a cysteine containing peptide can undergo oxidation to form intermolecular and intramolecular disulfide linkages resulting in cross-linked peptide antigens. **(c)** High-performance liquid chromatography chromatograms displaying extent of M27 peptide antigen disulfide cross-linking at 0, 0.5 and 48 hours in solution. Percent cross-linked is plotted over time as a function of cross-linked area-under-the-curve (AUC) divided by total AUC. **(d)** Frequency of cysteine and methionine in the human proteome (pie chart). Frequency of multiple cysteine residues in 25-mer peptides from the human proteome (rings). **(e)** Structural basis for isosteric substitution of oxidation prone WT amino acids with OXR amino acids. Abbreviations: APC = antigen presenting cell; WT = wild-type; OXR = oxidation resistant.

Humans have an estimated 10 billion different T cell clonotypes in circulation^6^, and each clone has a unique T cell receptor (TCR) that can bind to specific peptide-MHC complexes. Importantly, the ability of T cells to recognize different peptide-MHC complexes is key to the immune system’s ability to discern self from threats including cancer or infectious organisms. As T cells exist in a naïve state at low numbers prior to encountering their cognate antigen as a peptide-MHC complex, T cells must be activated, expanded, and differentiated to mount a response against a threat (e.g., pathogen or cancer)^7,8^, or suppress inflammation in the setting of autoimmunity^9^. To generate a T cell response in humans and other mammals, peptide-based vaccines utilize peptide antigens formulated in particles^10^ together with immunomodulators^11^. The particle promotes uptake by antigen-presenting cells (APCs), which process and present the peptide antigen as a peptide-MHC complex that is recognized by the TCR of specific T cell clones, whereas the immunomodulator helps to promote T cell activation and differentiation.

While peptide-based vaccines have shown promise for a variety of applications, amino acid sequence variability gives rise to differences in peptide antigen physical and chemical properties that present challenges to manufacturing and product stability^12^. For example, peptide antigens with the sulfur containing amino acids cysteine and methionine are prone to undergo oxidation resulting in impurities that can reduce shelf-life and impact biological activity. While the sulfhydryl group of cysteine is readily oxidized to yield disulfide linkages that are key structural features of many proteins, cysteines present in peptide antigens often undergo unwanted intra- or inter-molecular disulfide bonds (**Fig. 1b**), or the cysteine directly reacts with oxygen to form sulfenic, sulfinic or cysteic acid impurities^13^. Similarly, the sulfur atom in methionine is susceptible to oxidation, yielding methionine sulfoxide (MetO) as well as methionine sulfone^14^. These reactions and their byproducts result in impurities that impact product stability, shelf-life and can also interfere with MHC binding or T cell recognition^15,16^.

A variety of strategies have been developed to avoid or limit the impact of cysteine and methionine oxidation. Peptide antigens for use in vaccines can be selected to avoid those rich in cysteines and methionine, though, this may not be practical for pathogens rich in cysteines and methionine, such as certain viruses (e.g., HPV)^17^ and could unnecessarily limit the effectiveness of the vaccine. Other strategies are to include excipients in the vaccine formulation that function as reducing agents to prevent or reverse oxidation^18-20^. However, some excipient reducing agents can result in unwanted reactions or require additional testing in animals and humans and are often not 100% effective. Vialing the vaccine under inert gases (e.g., nitrogen), can also reduce oxidation but dramatically increases manufacturing costs and complexity and requires stability at all points up until vialing^21^. A potentially promising alternative strategy is to substitute sulfur-containing cysteine and methionine residues of peptide antigens with oxidation resistant (OXR) isosteres that are structurally related but omit sulfur.

The potential to substitute cysteine and methionine with OXR isosteres is supported by the observation that a single TCR can recognize multiple related peptide-MHC complexes with similar binding affinity^22^ and that conservative amino acid substitutions to peptide antigens are permissible^23^. For example, peptide-based vaccines used in preclinical studies of melanoma in mice commonly use a modified peptide antigen (TAPDNLGY<u>M</u>) that includes a methionine-substitution in place of an alanine present on the WT antigen (TAPDNLGY<u>A</u>) naturally present on tumor cells^24^. The T cells induced with vaccines containing the modified antigen also recognize the WT antigen and mediate anti-tumor efficacy. In this and similar instances, amino acids of peptide antigens have been substituted to improve binding to MHC molecules^25, 26^. While these examples highlight certain instances where amino acid substitutions are permissible, there are innumerable cases where a single amino acid change in a peptide antigen results in a complete loss of MHC binding or T cell recognition of the peptide-MHC complex^24^, and to our knowledge, a generalizable approach for replacing sulfur containing amino acids in peptide antigens has yet to be described. Therefore, there remains an unmet need for the identification of suitable substitutes for cysteine and methionine that are oxidation resistant.

To address this need, we sought to identify isosteric amino acids for cysteine and methionine that eliminate risks of oxidation, while retaining desired biological activity. We focused on the relatively narrow subset of possible isosteric amino acids that were oxidation resistant and similar in structure to cysteine and methionine. Herein, we show that the cysteine and methionine residues of peptide antigens substituted with the OXR amino acids alpha-aminobutyric acid and norleucine, respectively, are as effective as WT peptide antigens for inducing T cell immunity when utilized in peptide-based vaccines.

## Results

### Impact of cysteine oxidation on peptide antigen stability

While oxidation is appreciated as a commonly encountered problem during peptide antigen manufacturing, it is not well characterized in the literature. Therefore, our initial studies assessed how the presence of a single cysteine can impact the stability of a model peptide-based vaccine formulation during manufacturing and storage. As a model system, we quantified the kinetics of cysteine oxidation for a cysteine-containing peptide antigen (“M27”) in solution under ambient air. Notably >50% of M27 underwent oxidation over 48 hours at room temperature as indicated by a shift in retention time when assessed by high-performance liquid chromatography (HPLC) (**Fig. 1c**), which is commonly used to assess peptide antigen purity. We further characterized this peptide retention time shift with mass spectrometry and reaction with the reducing agent TCEP to demonstrate that the change is primarily attributable to cysteine disulfide formation (**Fig. 1c and Supplementary Fig. S1**). Such a marked change in purity of the peptide antigen due to disulfide formation can present challenges to regulated products used clinically.

The full scope of this problem is evident when considering that up to 43.9% of 25-mer peptide antigens generated from protein-coding DNA (relevant to peptide-based vaccines for cancer and autoimmunity) contain one or more cysteine residues and 11% of 25-mer peptides contain two cysteine residues (**Fig. 1d, Supplementary Calculation 1**) as calculated from the binomial distribution frequency with human proteome cysteine frequency of 2.3%^27^. Given the common occurrence of oxidation prone residues in peptide antigens there remains a need to reduce the occurrence of oxidation while maintaining epitope integrity for MHC binding and T cell recognition.

### Selection of oxidation resistant isosteres for cysteine and methionine

We next sought to investigate suitable substitutes for cysteine and methionine that are oxidation resistant (OXR) but are immunologically equivalent to the wild type (WT) antigen. The R group of cysteine comprises a methylene group linked to a sulfhydryl (-CH_2_-SH) group (**Fig. 1e**). One possible isostere of cysteine is serine (R = –CH_2_–OH), which replaces the sulfur with oxygen, a conservative change that has been demonstrated to be suitable for preserving protein structure^28^. However, as oxygen (Pauling electronegativity = 3.44) is more electronegative than sulfur (EN = 2.58), the polarity of the O–H bond is higher than the S–H bond, allowing the -OH group to readily donate and accept hydrogen bonds. An alternative isostere of cysteine would be to replace the sulfhydryl group with a methyl group, yielding the non-natural amino acid alpha-aminobutyric acid (hereafter, aBut; R = –CH_2_–CH_3_). Carbon’s electronegativity is 2.55, and thus, carbon shares electrons with hydrogen in a similar polarity as sulfur.

Thus, aBut could function as a substitute for cysteine that is not prone to oxidation. The R group of methionine comprises an ethylene group linked to a methyl group by a sulfur atom (–CH_2_–CH_2_–S–CH_3_). We examined 3 possible isosteres of methionine: leucine, norvaline, and norleucine (**Fig. 1e**). Leucine has a four carbon R-group, arranged with a branch point (R = –CH_2_– CH(CH_3_)_2_). Norvaline has a three-carbon linear R-group (R = –CH_2_–CH_2_–CH_3_). Replacing the sulfur group of methionine with a methylene group (–CH_2_–) yields the non-natural amino acid norleucine (sometimes abbreviated nor; R = –CH_2_–CH_2_–CH_2_–CH_3_). Replacement of methionine with norleucine in certain recombinant proteins retain their function^29^.

Whether cysteine and methionine substitution with OXR amino acids are permissible for use in peptide-based vaccines to generate immunity against WT antigens has not been previously explored. To address this, we formulated different peptide antigens containing either WT peptide antigens containing cysteine or methionine or the OXR versions as self-assembling nanoparticles (referred to as “SNAPvax”)^12^ and assessed whether OXR peptide antigens have similar immunogenicity to the WT peptide antigens (**Fig. 2a**). Importantly, SNAPvax forms particles with similar size and net charge independent of the underlying peptide antigen sequence, which eliminates any physical or chemical differences between the OXR and WT peptide antigens that can affect pharmacokinetics and pharmacodynamics. Thus, any differences in immune responses are assumed to be directly due to the amino acid substitutions and not differences in peptide antigen delivery.

**Fig. 2:**
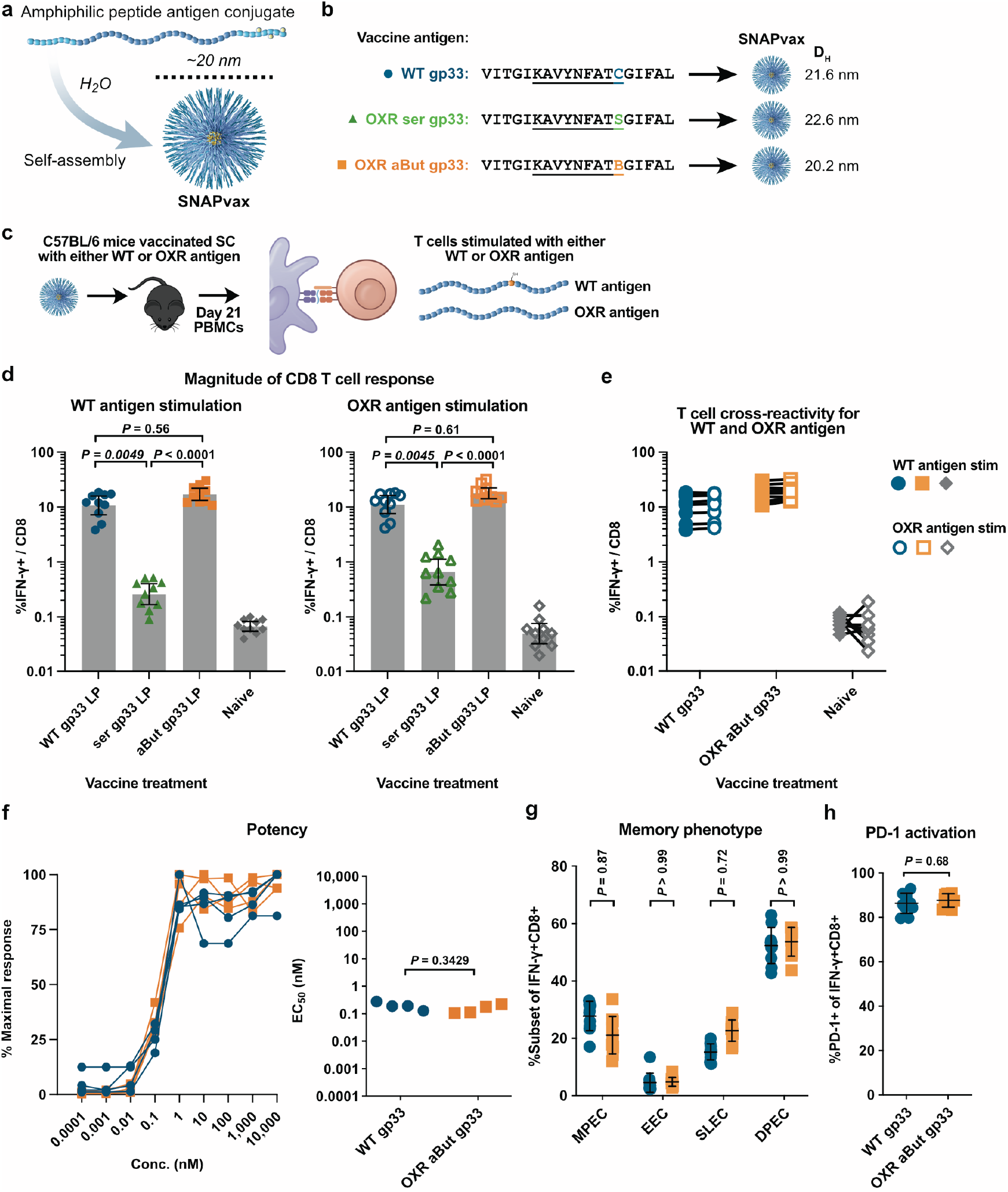
Replacing cysteine with aBut preserves T cell responses to viral antigen gp33. **(a)** Schematic showing SNAPvax self-assembly into ∼20 nm particles. **(b)** WT and OXR sequences of the viral antigen gp33. The 9-mer epitope known to bind MHC-I allele H-2D^b^ is underlined. The cysteine in WT gp33 (blue C) is substituted with alpha-aminobutyric acid in OXR aBut gp33 (orange B) or serine in OXR ser gp33 (green S). The cysteine is known to function as an MHC anchor residue. Each peptide was formulated as SNAPvax; hydrodynamic diameter as measured by dynamic light scattering is shown. (**c-h**) C57BL/6 mice (n = 10 per group) were immunized on day 0 and day 14 and assessed for T cell responses from whole blood on day 21 and spleen on day 34. **(c)** Cartoon schematic of experimental design. **(d)** Antigen-specific (IFN-γ+) CD8 T cell responses to either WT gp33 (left) or OXR aBut gp33 (right) stimulation. *P*-values assessed by one-way ANOVA (Kruskal-Wallis with Dunn’s correction). Error bars are geometric mean with 95% CI. **(e)** Data from (d) comparing WT vs OXR T cell response for each group. **(f)** Splenocytes were harvested on day 34 and stimulated with a dilution series of WT gp33 for 6 hours and assessed for IFN-γ by intracellular cytokine staining. The y-axis is the % of the IFN-γ response at a given concentration as a percentage of the maximal response. The interpolated potency EC_50_ was assessed using a non-linear sigmoidal regression and is graphed for each mouse (right). *P*-value assessed by Mann-Whitney. **(g)** Memory phenotype of IFN-γ+ cells on day 21 following WT gp33 stimulation. Bar graph shows quantification of individual mouse response for SLEC, EEC, MPEC and DPEC populations. P-values are by two-way ANOVA (Sidak’s multiple comparison test) comparing responses between groups. Error bars are mean ± standard deviation. **(h)** Activation status (PD-1 expression) of IFN-γ+ cells on day 21. *P*-value assessed by Mann-Whitney. Error bars are mean ± standard deviation. SLEC = short lived effector. EEC = early effector. MPEC = memory precursor. DPEC = double positive. D_h_ = hydrodynamic diameter.

### Assessment of suitable isosteres for cysteine

We first assessed the impact of isosteric substitution of cysteine at an anchor residue position. The immunodominant epitope of lymphocytic choriomeningitis virus (LCMV) gp33 (KAVYNFATC) contains a cysteine at the C-terminal anchor residue position for the relevant MHC Class I allele (MHC-I), H-2D^b^, of C57BL/6 mice and is a representative infectious disease antigen^30^. The C-terminal anchor residue position strongly influences binding to H-2D^b^, as substitution of methionine for cysteine at this location has been reported previously to improve affinity of the immunodominant gp33 epitope >250-fold^31^.

SNAPvax comprising the immunodominant epitope gp33 from LCMV were synthesized as the cysteine-containing WT or the OXR (aBut or serine) substituted versions. As expected, the different vaccines formed nanoparticles of similar diameter as assessed by dynamic light scattering (DLS) (**Fig. 2b**). Importantly, both the WT and OXR vaccines were freshly formulated and immediately used to avoid any impact of oxidation of the WT antigens on immunogenicity, which was confirmed by HPLC (**Supplementary Fig. 2**). Mice were treated by the SC route on days 0 and 14. The T cell response was assessed on day 7 and 21 by stimulating PBMCs with the WT, cysteine-containing minimal CD8 epitope (**Fig. 2c**). As expected, the WT treatment induced a high magnitude CD8 T cell response as assessed on day 7 (**Supplementary Fig. 2**) and day 21 (**Fig. 2d**). However, the serine substituted treatment induced WT antigen-specific T cells that were only slightly higher magnitude than the naïve group. This indicates that serine – even though the sulfur to oxygen substitution is chemically conservative and from the same group within the periodic table – is not a suitable isostere for cysteine.

In contrast, the aBut substituted treatment induced an equivalent magnitude CD8 T cell response as the WT treatment when cells from vaccinated animals were stimulated with the WT gp33 epitope (**Fig. 2d**). We further evaluated whether T cells induced with WT or OXR aBut treatments recognized WT gp33 peptide (solid shapes **Fig. 2e**) or OXR aBut gp33 peptide (open shapes **Fig. 2e**). The CD8 T cell response for both WT and OXR aBut treated animals were equivalent regardless of the peptide used for stimulation, demonstrating that T cells induced with OXR antigens cross-reacted with WT antigens and vice versa. Since the magnitudes were similar, we then undertook an indirect assessment of TCR potency for pMHC. Splenocytes from treated animals were stimulated with a dilution series of the WT gp33 peptide, and the percent of antigen-specific CD8 T cells producing IFN-γ was assessed by ICS and reported as a percentage of maximal response. There was no difference in the potency (EC_50_) measured for T cells from mice treated with either the WT gp33 or OXR aBut gp33 (*P* = 0.49; **Fig. 2f**) indicating that the WT and OXR antigens induced T cells with similar TCR potency for binding pMHC.

We further evaluated whether the T cell phenotypes resulting from vaccination with OXR aBut gp33 differed from that of WT gp33 vaccination. The memory phenotype – as assessed by KLRG1 and CD127 expression – of the WT gp33-specific CD8 T cells was the same whether vaccinated with WT gp33 or OXR aBut gp33 (**Fig. 2g**). Similarly, the activation status based on PD-1 expression was also the same (**Fig. 2h**). Similar magnitude and quality of immune responses between the WT and aBut OXR peptide antigens were also seen at day 7 indicating similar kinetics of responses as well (**Supplementary Fig. 2e-f**).

For an epitope with a TCR facing residue, we used the tumor-specific neoantigen M27 (LCPGNKYEM) derived from the murine tumor cell line B16-F10^32^ (**Fig. 3a**). In contrast to gp33, M27 contains a cysteine residue at position 2 of the MHC-I (H-2D^b^) epitope, which is not expected to act as an anchor residue typically found at either position 5 or 9^13^.

**Fig. 3:**
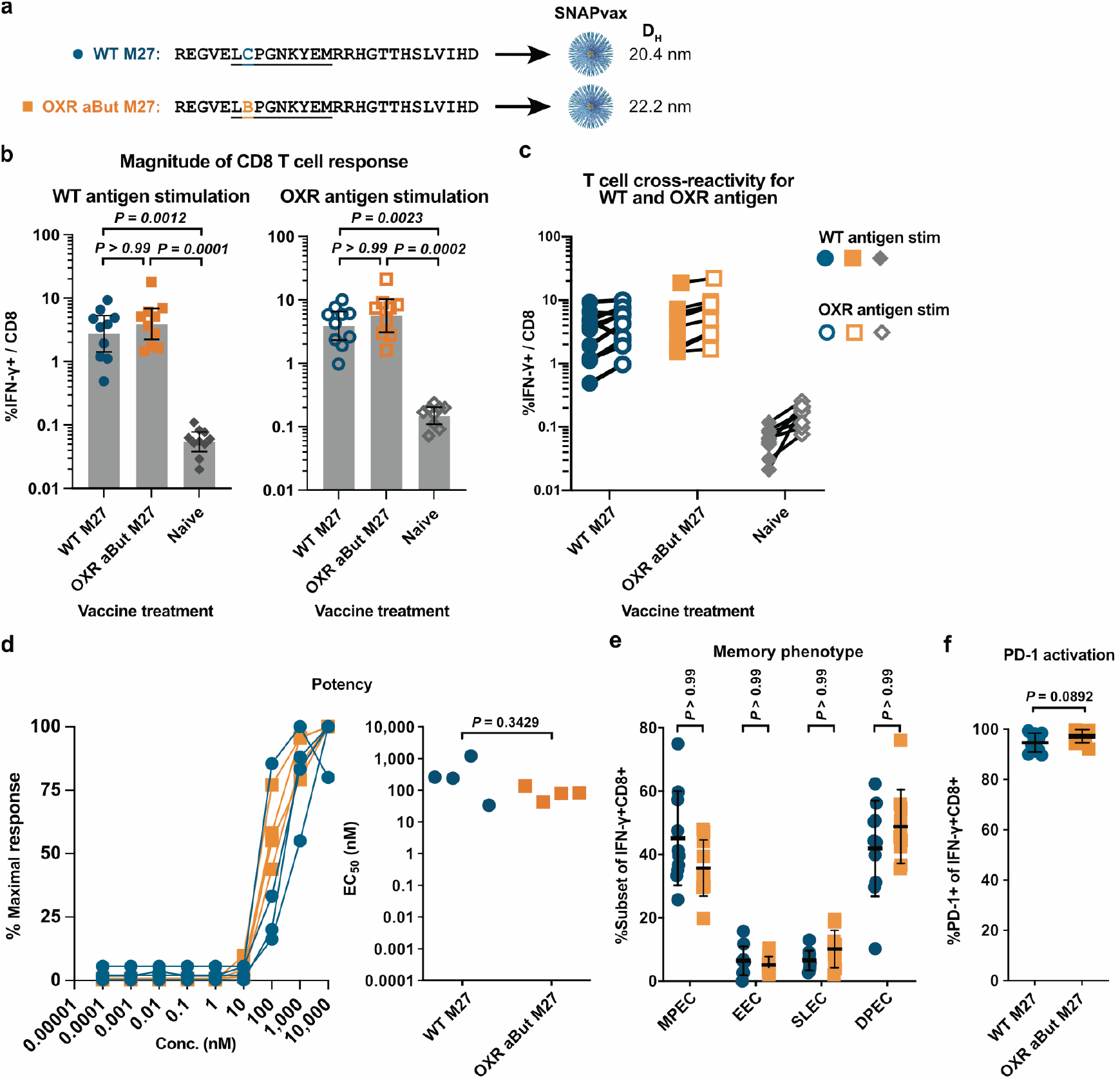
Alpha-aminobutyric acid substituted M27 is immunologically equivalent to wild type M27. **(a)** WT and OXR sequences of the neoantigen M27. The 11-mer epitope known to bind MHC-I allele H-2D^b^ is underlined. The cysteine in M27 (blue C) is substituted with alpha-aminobutyric acid in aBut M27 (orange B). Each peptide was formulated as SNAPvax and formed uniform nanoparticles; D_h_ as measured by dynamic light scattering is shown. (**b-f**) C57BL/6 mice (n = 10 per group) were immunized on day 0 and day 14 and assessed for T cell response from whole blood on day 21 and spleen on day 34. (b) Antigen-specific (IFN-γ+) CD8 T cell responses from whole blood on day 21 as measured by intracellular cytokine staining following peptide stimulation with either WT M27 (left) or OXR aBut M27 (right). *P*-values assessed by one-way ANOVA (Kruskal-Wallis with Dunn’s correction). Error bars are geometric mean with 95% CI. **(c)** CD8 T cells from whole blood were stimulated with either WT M27 (solid) or OXR aBut M27 (open) peptides. **(d)** Splenocytes were harvested on day 34 and stimulated with a dilution series of WT M27 and assessed for IFN-γ by intracellular cytokine staining. The y-axis is the % of the IFN-γ response at a given concentration as a percentage of the maximal response. The interpolated potency EC_50_ was assessed using a non-linear sigmoidal regression and is graphed for each mouse (right). *P*-value assessed by Mann-Whitney. **(e)** Memory phenotype of IFN-γ+ cells on day 21 following WT M27 stimulation. Bar graph shows quantification of individual mouse splenocyte response for SLEC, EEC, MPEC and DPEC populations. P-values are by two-way ANOVA (Sidak’s multiple comparison test) comparing responses between groups. Error bars are mean ± standard deviation. **(f)** Activation status (PD-1 expression) of IFN-γ+ cells on day 21. *P*-value assessed by Mann-Whitney. Error bars are mean ± standard deviation SLEC = short lived effector. EEC = early effector. MPEC = memory precursor. DPEC = double positive. D_h_ = hydrodynamic diameter.

Both the WT and OXR versions of the M27 peptide antigen were formulated as SNAPvax and characterized by HPLC (**Supplementary Fig. 3**), which showed that the WT M27 conjugate underwent cross-linking at room temperature in solution whereas the OXR version did not. Mice were treated with either the native M27 or OXR aBut M27, prepared immediately prior to use, on days 0 and 14, and immunogenicity assessed on days 7 and 21. Consistent with the findings with gp33, animals treated with either the WT or OXR versions had similar magnitude CD8 T cell responses generated to both the WT and OXR M27 epitopes (**Fig. 3b & c**), and there was no difference in the relative potency (as assessed by EC_50_) of the T cells (*P* = 0.1143; **Fig. 3d**). T cells with either vaccine (WT or OXR) also induced similar memory phenotype (**Fig. 3e**) and activation status (**Fig. 3f**).

### Assessment of suitable isosteres for methionine

To evaluate efficacy of isosteric substitution of methionine, we evaluated the immune responses to two MHC class I epitopes in C57BL/6 mice containing methionine residues either as anchor and non-anchor residues in the MC38 tumor epitope Adpgk (ASMTNMELM) or as an anchor residue in the B16 tumor epitope Trp1 (TAPDNLGYM).

We initially evaluated T cell responses from mice vaccinated with Adpgk LP substituted with either norleucine, norvaline or leucine at each of the three methionine positions (**Supplementary Fig. 4**). While substitution with norleucine, norvaline and leucine all gave detectable CD8 T cell responses as assessed by IFN-γ intracellular cytokine staining (left panel, **Supplementary Fig. 4d**) and ASMTNMELM:H-2D^b^-PE tetramer staining (right panel, **Supplementary Fig. 4e**), substitution with norleucine led to the highest T cell responses against the WT Adpgk min epitope (∼10% antigen-specific of total CD8 T cells) that were about 4-10-fold higher than those induced with the norvaline or leucine substituted antigens . From these data, we decided to further evaluate norleucine as an isostere for methionine.

**Fig. 4:**
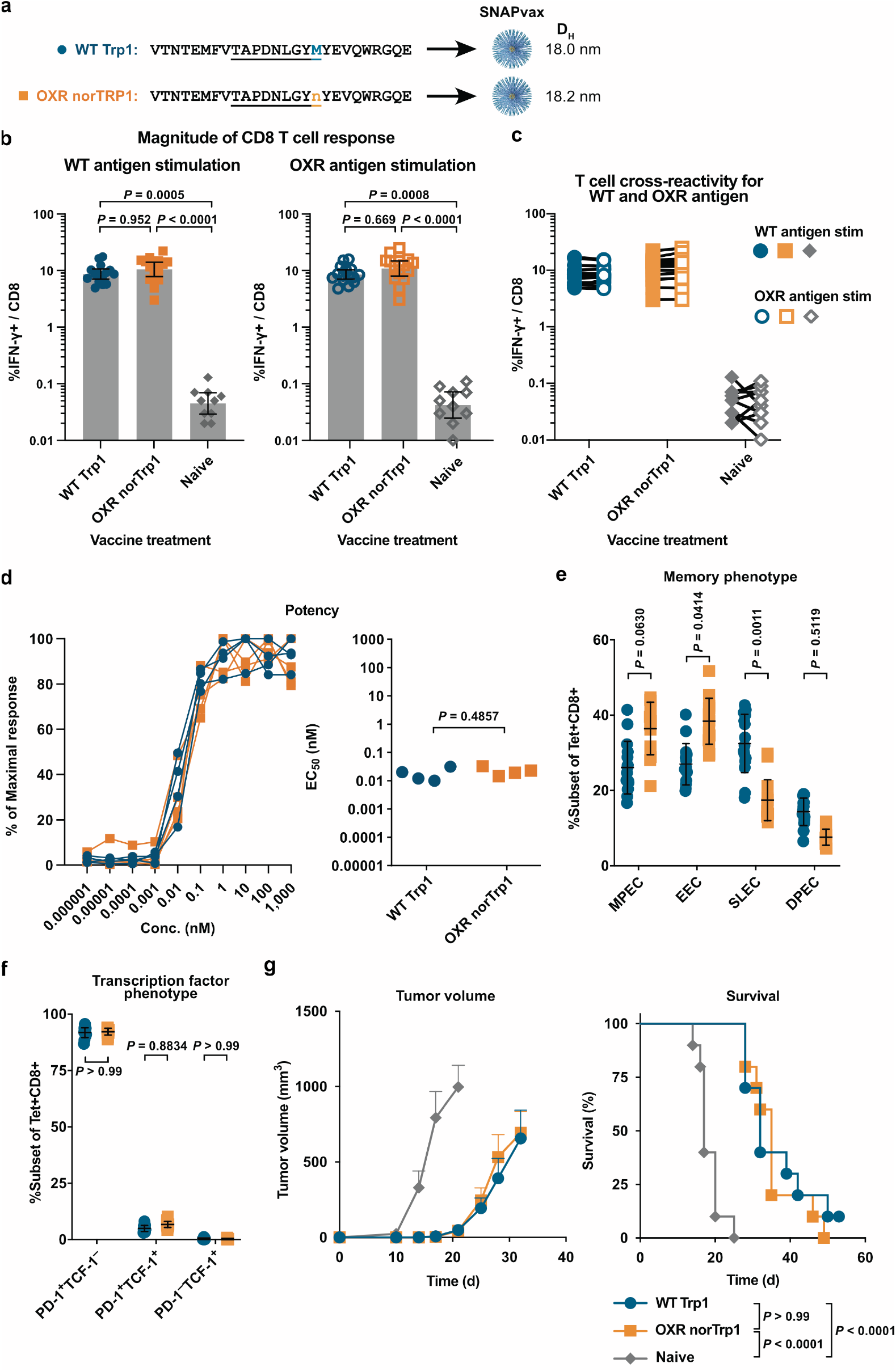
Norleucine substitution of methionine in Trp1 is immunologically equivalent to wild type Trp1. **(a)** Sequence of tumor self-antigen Trp1 and norleucine-substituted Trp1 (OXR norTrp1). The 9-mer minimal epitope known to bind MHC-I allele H-2D^b^ is underlined. The methionine in Trp1 (blue M) is substituted with norleucine in norTrp1 (orange n) and is known to function as an MHC anchor residue. Each peptide was formulated as SNAPvax and formed uniform nanoparticles; D_h_ as measured by dynamic light scattering is shown. **(b-g**) C57BL/6 mice (n = 10 per group) were immunized on day 7 and day 14 and assessed for T cell response from whole blood on day 21 and spleen on day 34. **(b)** Antigen-specific (IFN-γ+) CD8 T cell responses from whole blood on day 21 as measured by intracellular cytokine staining following peptide stimulation with either WT Trp1 (left) or OXR norTrp1 (right). *P*-values assessed by one-way ANOVA (Kruskal-Wallis with Dunn’s correction). Error bars are geometric mean with 95% CI. **(c)** CD8 T cells from whole blood were stimulated with either WT Trp1 (solid) or OXR norTrp1 (open) peptides. **(d)** Splenocytes were harvested on day 34 and stimulated with a dilution series of WT Trp1 and assessed for IFN-γ by intracellular cytokine staining. The y-axis is the % of the IFN-γ response at a given concentration as a percentage of the maximal response. The interpolated potency EC_50_ was assessed using a non-linear sigmoidal regression and is graphed for each mouse (right). *P*-value assessed by Mann-Whitney. **(e)** Memory phenotype of Tetramer+ cells on day 21. Bar graph shows quantification of individual mouse splenocyte response for SLEC, EEC, MPEC and DPEC populations. P-values are by two-way ANOVA (Sidak’s multiple comparison test) comparing responses between groups. Error bars are mean ± standard deviation. **(f)** Transcription factor phenotype and activation status (TCF-1 and PD-1 expression) of tetramer+ cells on day 21. P-values are by two-way ANOVA (Sidak’s multiple comparison test) comparing responses between groups. Error bars are mean ± standard deviation. **(g)** Tumor growth curves (left) and survival (right) following tumor challenge. Error bars are SEM. *P*-values for survival curves assessed by log-rank test with correction for multiple comparisons.

Trp1 is a CD8 epitope from the melanoma antigen tyrosinase related protein 1 (455-463) with A463M P9 anchor residue substitution demonstrated to improve binding to H-2D^b^ MHC Class I^33,34^. Herein, we refer to the methionine substituted epitope TAPDNLGYM as the WT epitope. Formulations of SNAPvax comprising the WT Trp1 or OXR norleucine substituted Trp1 (norTrp1) formed similar size particles (**Fig. 4a**) of high purity by HPLC (**Supplementary Fig. 5b**). To evaluate immunogenicity, C57BL/6 mice were vaccinated with SNAPvax comprising the Trp1 antigens on days 0 and 14 and T cell responses from PBMCs were evaluated on days 7 and 21. T cell function was also evaluated by tumor challenge using B16-F10 cells implanted at day 28.

**Fig. 5:**
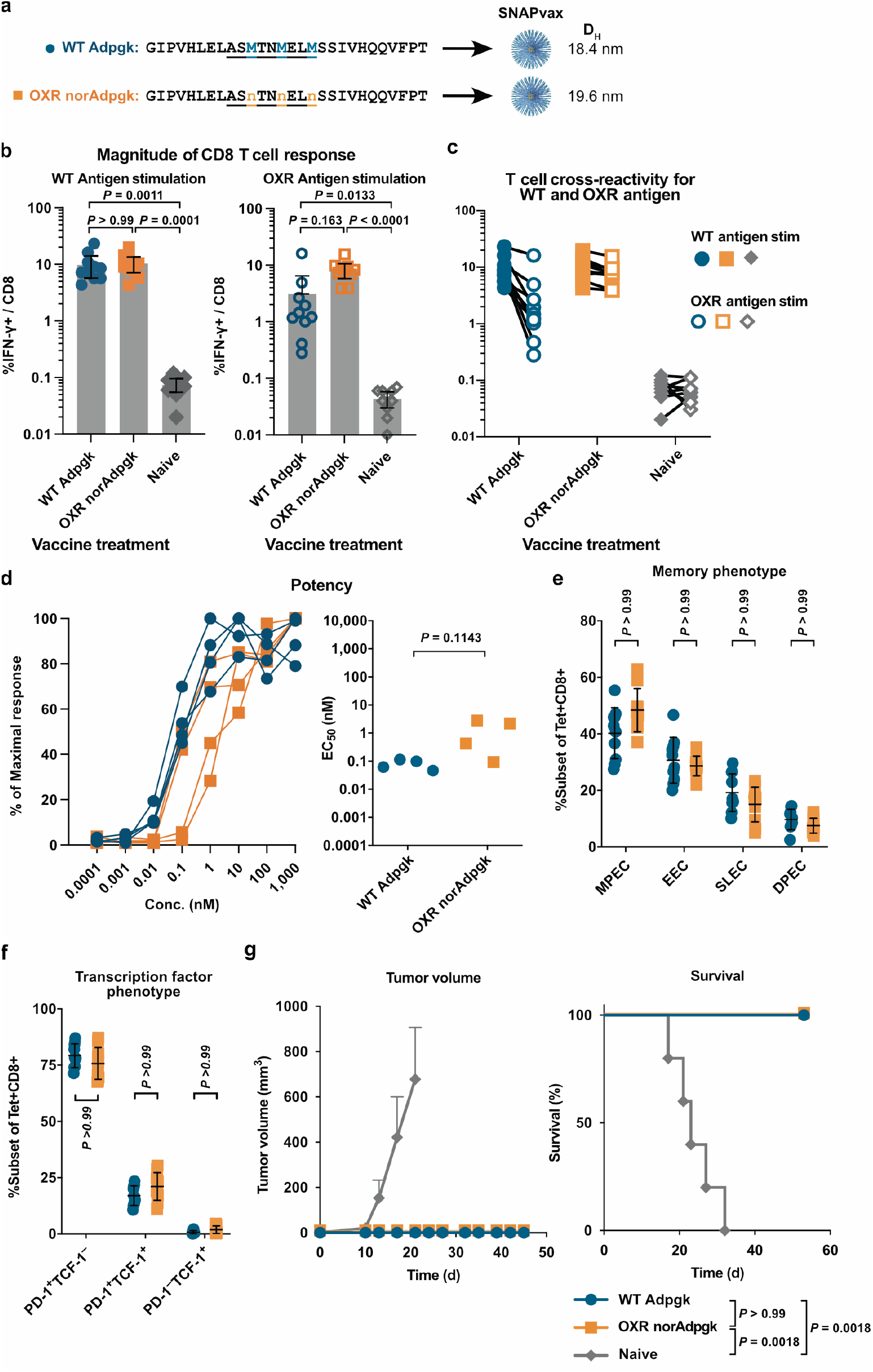
Norleucine substituted Adpgk is equivalent to wild type Adpgk. **(a)** Sequence of tumor self-antigen Adpgk and norleucine-substituted Adpgk (OXR norAdpgk). The 9-mer epitope is known to bind MHC-I allele H-2D^b^ and is underlined. The three Methionine residues in Adpgk (blue M) are substituted with norleucine in OXR norAdpgk (orange n). Each peptide was formulated as SNAPvax and formed uniform nanoparticles; D_h_ as measured by dynamic light scattering is shown. (**b-g**) C57BL/6 mice (n = 10 per group) were immunized on day 0 and day 14 and assessed for T cell response from whole blood on day 21 and spleen on day 34. **(b)** Antigen-specific (IFN-γ+) CD8 T cell responses from whole blood on day 21 as measured by intracellular cytokine staining following peptide stimulation with either WT Adpgk (left) or OXR norAdpgk (right). *P*-values assessed by one-way ANOVA (Kruskal-Wallis with Dunn’s correction). Error bars are geometric mean with 95% CI. **(c)** CD8 T cells from whole blood were stimulated with either WT Adpgk (solid) or OXR norAdpgk (open) peptides. **(d)** Splenocytes were harvested and stimulated with a dilution series of WT Adpgk and assessed for IFN-γ by intracellular cytokine staining. The y-axis is the % of the IFN-γ response at a given concentration as a percentage of the maximal response. The interpolated potency EC_50_ was assessed using a non-linear sigmoidal regression and is graphed for each mouse (right). *P*-value assessed by Mann-Whitney. **(e)** Memory phenotype of tetramer+ cells on day 21. Bar graph shows quantification of individual mouse response for SLEC, EEC, MPEC and DPEC populations. P-values are by two-way ANOVA (Sidak’s multiple comparison test) comparing responses between groups. Error bars are mean ± standard deviation. **(f)** Activation status (PD-1 expression) of tetramer+ cells on day 21. P-values are by two-way ANOVA (Sidak’s multiple comparison test) comparing responses between groups. Error bars are mean ± standard deviation. **(g)** Tumor growth curves (left) and survival (right) following tumor challenge. Error bars are SEM. *P*-values assessed by log-rank test with correction for multiple comparisons.

Mice treated with WT Trp1 and OXR norTrp1 had similar magnitude T cell responses to both the WT and OXR epitopes (**Fig. 4b & c**). There was also no difference in the T cell potency (EC_50_) for the WT Trp1 epitope (*P* = 0.4857; **Fig. 4d**). Furthermore, we evaluated T cell recognition by WT Trp1 tetramer staining, confirming that OXR norTrp1 vaccinated T cells recognize the WT Trp1 in the context of H-2D^b^ (**Supplementary Fig. 5d**).

Next, we assessed whether the T cell phenotypes resulting from vaccination with OXR norTrp1 differed from that of the WT Trp1. The memory phenotype – as assessed by KLRG1 and CD127 expression revealed that T cells from OXR norTrp1 vaccinated mice had a higher early effector population and lower short lived effector population that was statistically significant (P = 0.0414 and P = 0.0011 respectively) (**Fig. 4e**). Characterization of PD-1 and TCF-1 expression on tetramer positive CD8 cells, however, was equivalent between WT Trp1 and OXR norTrp1 vaccinated mice (**Fig. 4f**). Characterization of T cells from day 7 demonstrated similar trends. However, no significant differences in early effector or short-lived effector population were measured (**Supplementary Fig. 5g**).

To evaluate T cell function, mice were challenged with the B16-F10 tumor cell line at day 28 and were administered anti-PD-L1 twice on days 35 and 42. Mice vaccinated with WT Trp1 or OXR norTrp1 had improved tumor control compared to naïve mice (P < 0.0001) that were equivalent by both tumor growth and mean survival time (P > 0.99). (**Fig. 4g-h**)

The murine neoantigen Adpgk (ASMTNMELM) from the MC38 tumor cell line is an MHC class-I epitope for H-2D^b^ with 2 nM binding affinity^35^. The Adpgk epitope contains 3 methionine residues in both anchor (P9) and solvent exposed positions (P3, P6) including the mutated residue R303M. Formulations of SNAPvax comprising the WT Adpgk or norleucine substituted Adpgk (OXR norAdpgk) formed similar size particles (**Fig. 5a**) of high purity by HPLC (**Supplementary Fig. 6b**). To evaluate immunogenicity, C57BL/6 mice were vaccinated with SNAPvax formulations comprising WT Adpgk or OXR norAdpgk with T cell responses from PBMCs evaluated on days 7 and 21. T cell functionality was further evaluated by tumor challenge at day 28.

Mice treated with WT Adpgk had similar magnitude CD8 T cell responses to both the WT and OXR epitopes (**Fig. 5b & c**). The potency (EC_50_) for OXR T cells from the norAdpgk treated mice were on average higher than WT Adpgk treated mice, but the difference was not significant (P = 0.1143, n = 4) (**Fig. 5d**). Tetramer staining revealed a similar magnitude CD8 T cell response induced by the WT and OXR vaccines and that the T cell responses recognized the Adpgk epitope within the context of H-2D^b^ (**Supplementary Fig. 6e**). There were also no significant differences in activation or memory status of the T cells induced (**Fig. 5e & d**). Both WT Adpgk and OXR norAdpgk vaccinated mice also showed improved tumor control compared to naïve mice treated with anti-PD-L1 alone by both tumor volume (P = 0.0018) and mean survival time (**Fig. 5g**), substantiating that both WT and OXR vaccines induce comparable magnitude, potency and function of T cells.

## Discussion

Oxidation of the sulfur-containing amino acids cysteine and methionine can complicate the manufacturing of peptide antigens used in vaccines and characterization of immune responses. Cysteine and methionine are prevalent in epitopes used as peptide antigens for vaccines against viruses, cancer (e.g., Tp53^36^ and KRAS G12C^37^) and autoimmunity. We addressed this problem with a simple, generalizable solution: replace cysteine and methionine with alpha-aminobutyric acid (aBut) and norleucine (nor), respectively, to generate oxidation resistant (OXR) peptide antigens that can elicit CD8 T cell responses to naturally occurring, wild type (WT), antigens. Across multiple H2-D^b^ restricted epitopes with substitutions spanning both anchor and TCR-facing residues, OXR peptide antigens elicited comparable magnitude, quality and function (as assessed by tumor challenge) of CD8 T cell responses as compared to WT antigens when used as vaccines.

A related approach has been to use so-called heteroclitic peptide antigens wherein amino acid substitutions are designed to (i) prime a T cell response cross-reactive with the WT antigen (e.g., gp100^38^, Trp1^24^) or (ii) enhance T cell recognition *in vitro* (e.g., cysteine substitutions in tyrosinase 243–251^39^ and p68^40^ that overcome oxidative linked issues). In many cases, the substitutions used have been evaluated case-by-case and not specifically meant to address the issue of cysteine and methionine oxidation. An exception is the work by Sachs *et al*. who showed that aBut can be used as a replacement for avoiding oxidation effects of cysteine in several HLA-A*02:01 restricted epitopes^15^. Notably, aBut modified peptides were solely used to stimulate pre-existing T cell responses to WT antigen *in vitro*. Here, we show that OXR substitution can be used to generate *de novo*, cross-reactive T cell responses to both OXR and WT epitopes *in vivo*.

Though avoiding or replacing oxidation sensitive residues has definitive advantages for manufacturing, oxidized antigens can sometimes be biologically relevant. Indeed, methionine sulfoxide reductase (MsrA) is downregulated across multiple human cancers^41^ and correlates with breast cancer grade^42^. Furthermore, immunopeptidomics has identified that oxidized peptides can sometimes serve as a source of cancer-restricted antigens^43^ and may be preferred for use in certain therapeutic vaccines. As such, the decision whether to replace cysteine and methionine with OXR amino acids should be made on a case-by-case basis and informed by an understanding of the epitopes relevant to the disease. Notably, prior work reports instances where aBut substitution for cysteine can reduce recognition for specific epitopes^15^. Although we did not observe this in our models, larger, allele-diverse panels are needed to estimate the frequency of exceptions to our approach.

In summary, this work demonstrates that peptide-based vaccines with isosteric substitution of the sulfur-containing amino acids cysteine and methionine with alpha-aminobutyric acid and norleucine, respectively, maintain immunogenicity while removing a major source of degradation. Crucially, we established that TCR recognition of these isosteres is fully cross-reactive: OXR epitopes are capable of inducing *de novo* T cell responses that recognize the native WT antigens, while also serving as stable reagents for screening and measuring pre-existing responses against WT targets. This dual utility implies that OXR peptides are suitable for both designing therapeutic vaccines and facilitating high-throughput discovery screens. We anticipate the greatest impact in multi-epitope vaccines and large-scale immunomonitoring, where standardizing the oxidative state can translate into shorter development times and more reliable manufacturing of clinical products.

## Methods

Methods and any associated references are available in the online version of the paper.

## Notes

This work was supported by the Intramural Research Program of the VRC, NIAID, NIH. The findings and conclusions in this report are those of the authors and do not necessarily reflect the views of the funding agency or collaborators.

## Acknowledgements

The authors acknowledge the contributions of: Carmelo Chiedi, Sienna Rush, Oscar Hernandez, Gloria Salbador, and Diana Scorpio of the VRC Translational Research Program, NIH; Gloria Chu, Danielle Thompson, and Patricia Darrah of the Cellular Immunology Section, VRC, NIH.

## Author contributions

A.S.I. and G.M.L. conceived of the project. A.S.I. designed the experiments. A.S.I., A.M.S., M.M., M.E., and N.G.P performed the experiments. A.S.I., C.M.G., D.R.W. wrote the manuscript; R.A.S. and G.M.L. revised the manuscript. All authors had complete access to all data, contributed to the analysis, and reviewed the manuscript.

## Data availability statement

The data that support the findings of this study are available from the corresponding author upon request.

## Online methods

### Animal protocols

Animal experiments were conducted at the Vaccine Research Center at the National Institutes of Health. Animal protocols underwent review and were approved by the respective Animal Care and Use Committee (ACUC) before the start of experiments. Animal experiments complied with the respective ethical guidelines as set by each ACUC.

### Animals

Female C57BL/6 (B6) mice were obtained from The Jackson Laboratory and maintained under specific-pathogen-free conditions. B6 mice were 8–12 weeks of age at the start of experiments. Animals were randomly assigned to either control or experimental groups.

### Vaccines

Native peptide antigens and modified peptide antigens are listed in Supplementary Table 1 and custom synthesized by Genscript using standard solid-phase peptide synthesis and purified (>90%) by high-performance liquid chromatography. Imidazoquinoline-based TLR-7/8 agonists and peptide antigen conjugates were synthesized by Barinthus Biotherapeutics North America as previously described^12^.

Conjugate vaccines were produced by linking peptide antigen containing a C-terminal azidolysine to a hydrophobic block oligo-TLR7/8 agonist using copper-free strain-promoted azide-alkyne cycloaddition click chemistry reaction.

### Vaccinations

Vaccines were prepared in sterile, endotoxin-free (<0.05 endotoxin units mL−1) PBS (Gibco). For mice, vaccines were administered subcutaneously in a volume of 50 μL in each hind footpad. Animals treated with checkpoint inhibitor, anti-PD-L1 (clone 10F.9G2, BioXCell catalog no. BE0101), received 200 μg administered by the intraperitoneal (i.p.) route in 100 μL of PBS.

### Tumor cell lines and implantation

B16-F10 was acquired from ATCC (CRL-6475). B16-Adpgk was derived from B16^5^. Working cell banks (passage 4) were generated immediately on receipt and used for tumor experiments. Cells were determined to be mycoplasma free on establishment of each working cell bank. B16-F10 and B16-Adpgk were cultured in RPMI-1640 media (GE Life Sciences) supplemented with 10% v/v heat-inactivated FCS (Atlanta Biologicals), 100 U mL^−1^ penicillin, 100 μg mL^−1^ streptomycin (Gibco), 1× nonessential amino acids (GE Life Sciences) and 1 mM sodium pyruvate (GE life Sciences). For each tumor cell implantation, a frozen aliquot was thawed, passaged once and gathered using trypsin EDTA (Gibco), quenched with HI-FCS, washed in PBS and implanted subcutaneously in sterile PBS. Tumors were measured using digital calipers twice per week, and tumor volume was estimated using the formula (tumor volume = short × short × long/2). Animals were euthanized when tumors reached size criteria (1,000 mm^3^).

### Sample processing

Measurement of antigen-specific CD8 T cell responses by intracellular cytokine staining (ICS) was performed as previously described^12^. Briefly, 200 μL of heparin-treated whole blood was RBC-lysed with ACK lysis buffer (Quality Biologicals), filtered, and cultured in complete RPMI in 96-well plates with 2.5 μg mL^−1^ of the specified stimulation peptide antigen (Genscript). Brefeldin A (BD) was added to a final concentration of 10 μg mL^−1^ 2 h after purified peptides were added and cells were incubated for an additional 4 h. After washing with PBS, cells were stained with ultraviolet Blue Live-Dead Dye (Life Technologies), washed and blocked with anti-CD16/CD32 (clone 2.4G2, BD catalog no. 553142) for 10 min at room temperature. After blocking, cells were surface stained for 30 min at room temperature. Cells were washed, suspended in BD Perm/Wash buffer containing intracellular stain for 30 min at 4°C. Cells were washed and suspended in 0.5% paraformaldehyde (Electron Microscopy Sciences) in PBS and then evaluated by flow cytometry. See **Supplementary Table 2** for antibodies used for surface and intracellular staining.

For tetramer staining, cells were isolated as for intracellular cytokine staining. Instead of peptide stimulation, cells were immediately stained with ultraviolet Blue Live-Dead Dye (Life Technologies) in the presence of dasatinib (Cat # S1021). Cells were stained with H-2D^b^-PE tetramers with specific peptides: Trp1:TAPDNLGYM and Adpgk:ASMTNMELM. Tetramers were produced as described previously^44^. Cells were then blocked and surface stained as for intracellular cytokine staining above, then permeabilized with eBioscience Foxp3 permeabilization buffer (Cat # 00-5523-00), stained overnight at 4°C, washed, and resuspended in 0.5% paraformaldehyde in PBS.

### Flow cytometry

Samples were acquired on a modified BD LSR Fortessa X-50 flow cytometer running BD FACSDiva software v.8.0.1.

### Statistical analyses

#### Data analysis

Flow cytometry data was analyzed using FlowJo v10. Statistical analyses were performed with Prism 10 (GraphPad). Graphs were rendered in FlowJo and Prism.

