## Supplementary Information for "Improving peptide vaccine manufacturability without sacrificing immunogenicity: substitution of methionine and cysteine with oxidation-resistant isosteres"

*

† Corresponding authors: David R. Wilson and Geoffrey M. Lynn


**Supplemental Materials.**

**Supplementary Calculation 1.** Cysteine frequency in peptide antigens
**Supplementary figure 1.** Oxidation reversal of M27 antigen using TCEP
**Supplementary figure 2.** alpha-aminobutyric acid substituted gp33 induces cross-reactive CD8 T cells
**Supplementary figure 3.** Oxidation effect on M27 cysteine residue
**Supplementary figure 4.** Isosteric substitution of methionine in Adpgk with norleucine, norvaline or leucine
**Supplementary figure 5.** Norleucine substitution of methionine in Trp1 is equivalent to native Trp1
**Supplementary figure 6.** Norleucine substituted Adpgk is equivalent to native Adpgk
**Supplementary Table 1.** Peptide antigens
**Supplementary Table 2**. Flow cytometry reagents

**Supplementary Calculation 1.** Cysteine frequency in peptide antigens

The frequency of cysteine residues in the human proteome varies depending on the source. For the primary figure calculation, we used the 2.3% cysteine frequency reported Wiedman *et. al.*[1] for all human proteins in UniProt, which is close to the 2.0% estimated by Sachs *et. al.*[2] The theoretical probability of finding a 25-mer peptide with a specific number of cysteine residues was calculated using a binomial distribution. The probability P(k) of a peptide of length n containing exactly k cysteine residues is given by the formula:

$$P(k) = \binom{n}{k} * p^k (1-p)^{n-k}$$

Where:

- n = 25 (the length of the peptide antigen)
- k = the number of cysteine residues (1, 2, or 3)
- p = 0.02 (the reference frequency of cysteine in the human proteome)
- $\binom{n}{k} = \frac{n!}{k!(n-k)!}$ is the binomial coefficient, representing the number of ways to choose k positions for cysteine out of n total positions. Binomial coefficient $\binom{n}{k}$ for 1, 2 or 3 cysteines in a 25-mer peptide was 25, 300 or 2300 respectively.

Individual frequencies of a 25-mer peptide antigen having one, two, three or four cysteine residues was calculated to be 32.9%, 9.3%, 1.7% and 0.039% respectively using a cysteine frequency in the proteome of 2.3%.

For the cumulative binomial probabilities of a 25-mer peptide to have ≥1, ≥2, or ≥3 cysteines we use the following equations.

$$P(x \geq k) = 1 - P(x < k) = 1 - \sum_{i=0}^{k-1} P(x = i)$$

Where,

$$P(x) = P(k) = \binom{n}{k} * p^k (1-p)^{n-k}$$

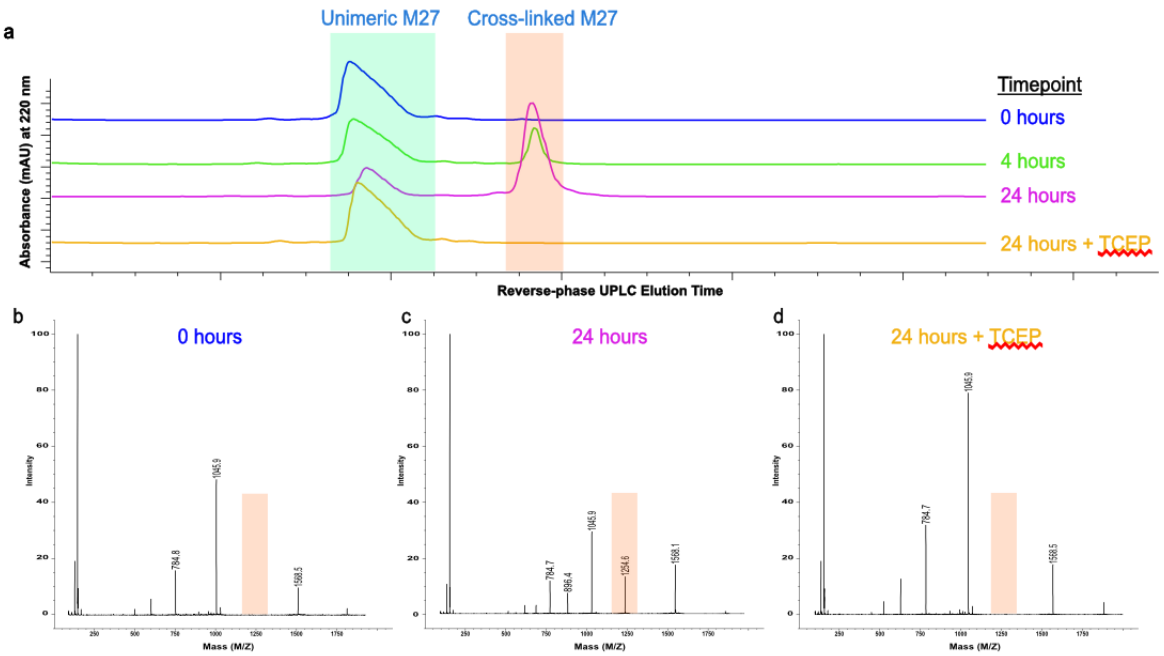


**Supplementary figure 1 | Oxidation reversal of M27 antigen using TCEP**

**(a)** Reverse phase UPLC traces using ESI-MS detection of WT M27 antigen over time with y-axis denoting absorbance at 220 nm in arbitrary absorbance units (mAU). The peaks highlighted in green shows the retention time of the unimer M27 peptide whereas the peak highlighted in orange shows the retention time of the crosslinked peptide absorbance.

**(a)** An averaged mass spectrum over the 0-hour UPLC trace from 4.5-5.3 minutes representative of the unoxidized, unimer antigen.

**(b)** An averaged mass spectrum over the 24-hour UPLC trace from 4.5-5.3 minutes representative of the oxidized, disulfide crosslinked, dimer antigen.

An averaged mass spectrum over the UPLC trace from 4.5-5.3 minutes after TCEP addition representative of the unoxidized, unimer antigen.

**Methods:** A solid sample of the WT M27 LP antigen (REGVELCPGNKYEMRRHGTTHSLVIHD, Genscript) was dissolved in DMSO (Gaylord) to a concentration of 10 mM, then diluted 1:10 in 1x PBS (Gibco) to yield a final solution of 1 mM antigen in 10% DMSO. Within 2 minutes of dilution, the sample was analyzed by UPLC-MS using a 20-40% ACN gradient over 10 minutes. The unoxidized unimer eluted at ~4.5 minutes, and ESI-MS confirmed the expected molecular weight [m/z = 3135.54, observed 1568.5 for $(M+2H)^{2+}$]. The sample was left at room temperature for 4 hours and re-analyzed. UPLC-MS revealed a new peak at 5.3 minutes, corresponding to ~30% disulfide crosslinking. This was confirmed by mass spectral peaks at m/z 896.4 and 1254.5, consistent with the dimerized species (MW = 6269.05) as $(M+7H)^{7+}$ and $(M+5H)^{5+}$ ions. After 24 hours, the oxidized fraction increased to 71%. To test reversibility, 20 molar equivalents of tris(2-carboxyethyl)phosphine (TCEP, Thermo Fisher Scientific) were added by solid and vortexed to ensure full mixture into the antigen solution. Subsequent UPLC-MS analysis showed complete reduction of the crosslinked dimer back to its unoxidized, unimer form. ESI-MS confirmed the reduction of this species by the loss of peaks in the mass spectrum representative of the oxidized dimer (Supplementary figure 1d).

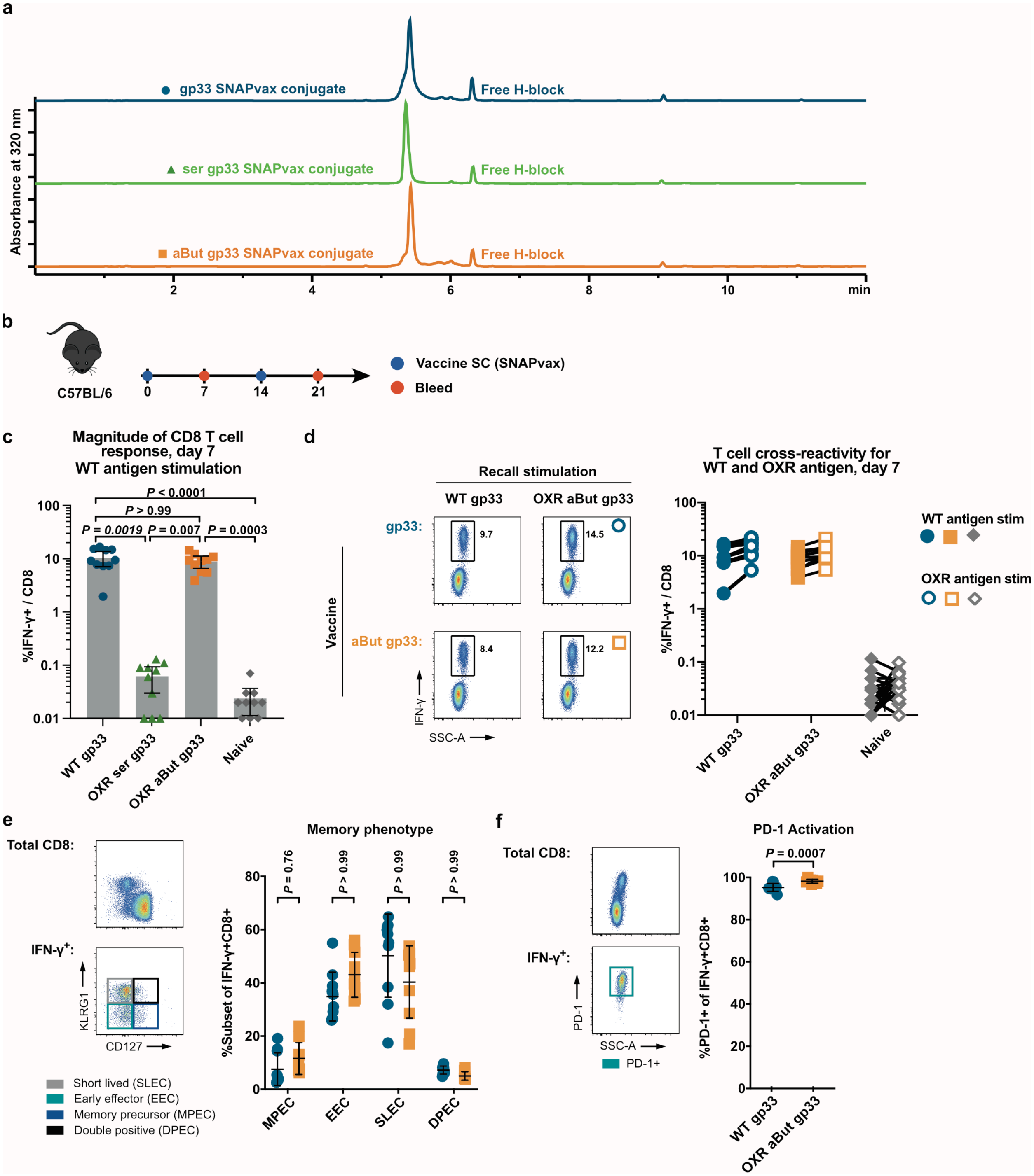


**Supplementary figure 2 | Alpha-aminobutyric acid substituted gp33 induces cross-reactive CD8 T cells**

- **(a)** Reverse phase HPLC traces of SNAPvax conjugates of WT gp33(blue), OXR aBut gp33 (orange), and ser gp33 LP (green) with y-axis showing absorbance at 320 nm in arbitrary absorbance units (mAU). "Free H-block" is a controlled component of the SNAPvax formulation and is not related to the peptide antigen.
- **(b)** C57BL/6 mice (*n* = 10 per group) were treated subcutaneously (SC) on day 0 and 14 and bled on day 7 and 21. Data presented in (c-f).

**(c)** Antigen-specific (IFN-γ+) CD8 T cell responses to WT gp33. *P*-values assessed by one-way ANOVA (Kruskal-Wallis with Dunn's correction). Error bars are geometric mean with 95% CI.

**(d)** CD8 T cell responses on day 7 using either WT gp33 min or aBut gp33 min peptide stimulation with IFN-γ staining as measured in (c).

**(e)** Memory phenotype of IFN-γ+ cells on day 7 following WT gp33 min stimulation. Bar graph shows quantification of individual mouse splenocyte response for SLEC, EEC, MPEC and DPEC populations. P-values are by two-way ANOVA (Sidak's multiple comparison test) comparing responses between groups. Error bars are mean ± standard deviation.

**(f)** Activation status (PD-1 expression) of IFN-γ+ cells on day 7. Histograms show concatenated samples of all acquired events for total CD8 T cells (top left) and IFN-γ+ CD8 T cells (bottom left), and bar graph shows quantification of each individual sample (right). *P*-value assessed by Mann-Whitney.

Abbreviations: WT = wildtype. OXR = oxidation resistant. aBut = aminobutyric acid. SLEC = short lived effector. EEC = early effector. MPEC = memory precursor. DPEC = double positive. $D_h$ = hydrodynamic diameter.

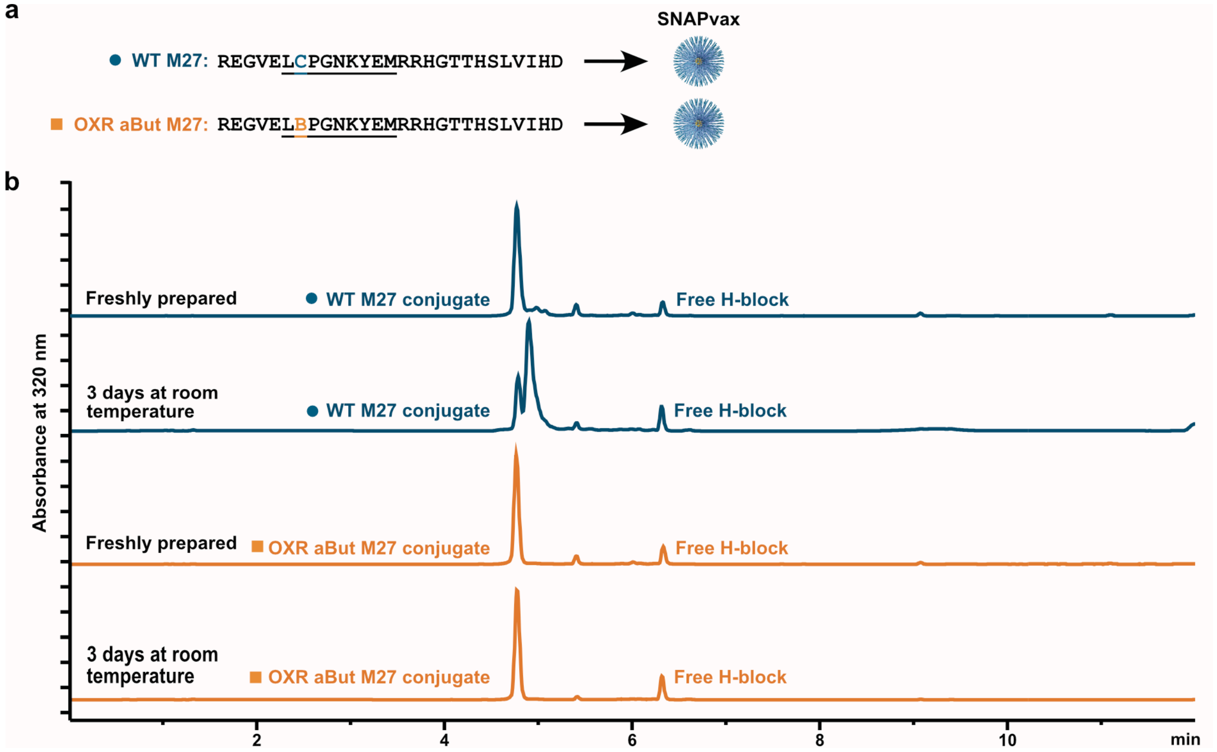


**Supplementary figure 3 | Oxidation effect on M27 cysteine residue**

**(a)** Sequence of the neoantigen M27. The 11-mer epitope known to bind MHC-I allele H-2D$^b$ is underlined and bold. The central cysteine in M27 (blue C) is substituted with butyrate in OXR aBut M27 (orange B). Each peptide was formulated as SNAPvax nanoparticles.

**(b)** Reverse phase HPLC traces of SNAPvax conjugates of WT M27 LP (blue), OXR aBut M27 (orange) either freshly prepared or after three days at room temperature as 10% DMSO in aqueous buffer. Y-axis shows absorbance at 320 nm in arbitrary absorbance units (mAU).

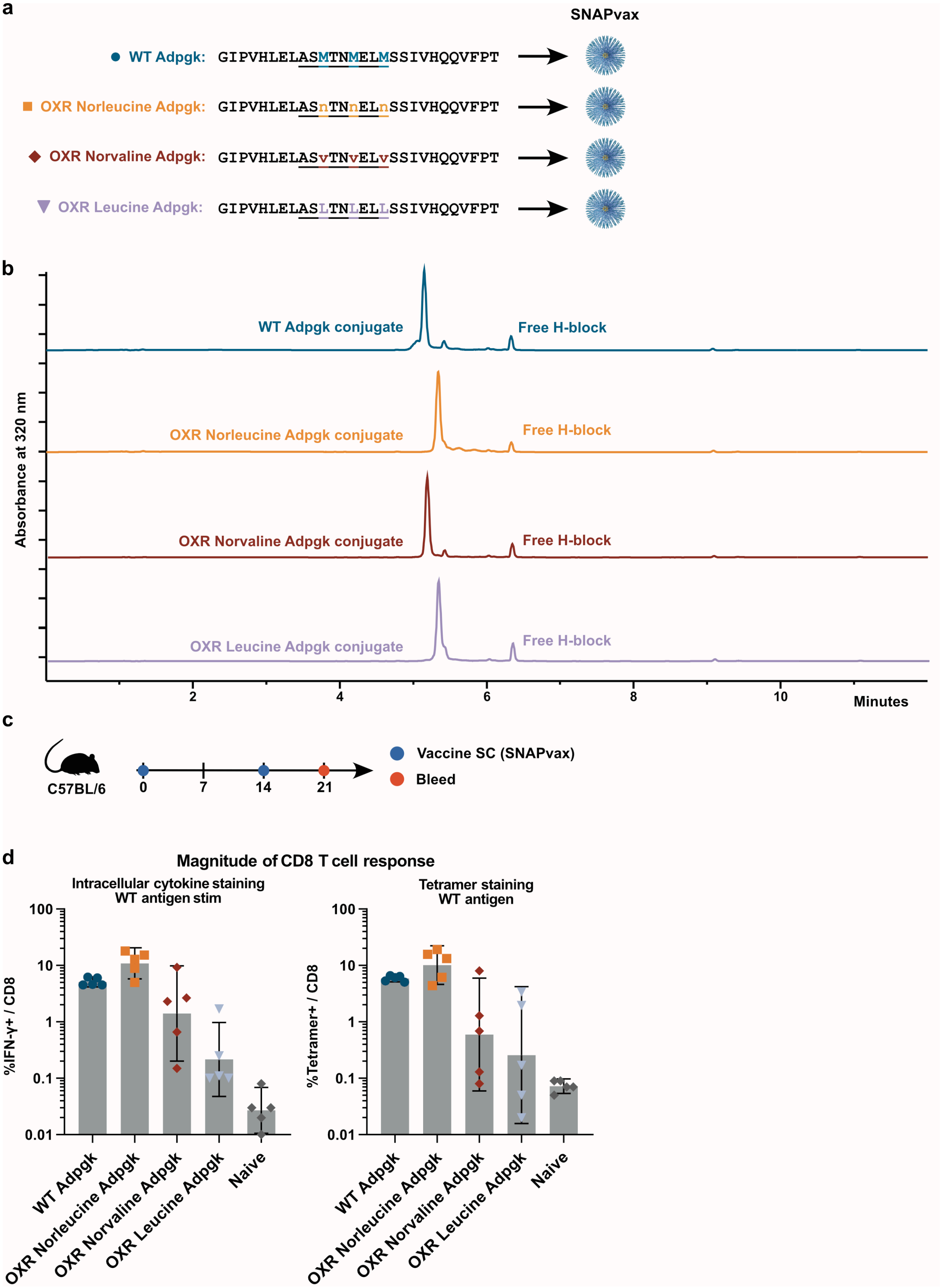
a
SNAPvax
WT Adpgk: GIPVHLELASMTNMELMSSIVHQQVFPT
OXR Norleucine Adpgk: GIPVHLELASnTNnELnSSIVHQQVFPT
OXR Norvaline Adpgk: GIPVHLELASvTNvELvSSIVHQQVFPT
OXR Leucine Adpgk: GIPVHLELASLTNLELLSSIVHQQVFPT
b
Absorbance at 320 nm
WT Adpgk conjugate
Free H-block
OXR Norleucine Adpgk conjugate
Free H-block
OXR Norvaline Adpgk conjugate
Free H-block
OXR Leucine Adpgk conjugate
Free H-block
2
4
6
8
10
Minutes
c
C57BL/6
0
7
14
21
Vaccine SC (SNAPvax)
Bleed
d
Magnitude of CD8 T cell response
Intracellular cytokine staining
WT antigen stim
%IFN-γ+ / CD8
Tetramer staining
WT antigen
%Tetramer+ / CD8
100
10
1
0.1
0.01
WT Adpgk
OXR Norleucine Adpgk
OXR Norvaline Adpgk
OXR Leucine Adpgk
Naive

**Supplementary figure 4 | Isosteric substitution of methionine in Adpgk with norleucine, norvaline or leucine**

**(a)** Sequence of WT Adpgk, norleucine substituted Adpgk (OXR norAdpgk), norvaline substituted Adpgk, and leucine substituted Adpgk. The 9-mer epitope known to bind MHC-I allele H-2D$^b$ is underlined and bold. The three methionine residues of Adpgk (blue M) are substituted with norleucine, norvaline or leucine respectively in the three isosteric peptides. Each peptide was formulated as SNAPvax and formed nanoparticles.

**(b)** Reverse phase HPLC traces of SNAPvax conjugates of WT Adpgk (blue) and norleucine Adpgk (orange), norvaline Adpgk (norvaline) or leucine Adpgk (purple) with y-axis showing absorbance at 320 nm in arbitrary absorbance units (mAU). The conjugates elute at similar times. As designed, an approximately 10% molar excess of free H-block is present in the formulation, which incorporates in the interior of the self-assembling nanoparticle micelle upon reconstitution in aqueous buffer.

**(c)** C57BL/6 mice (n = 5 per group) were vaccinated subcutaneously (SC) on day 0 and 14 and T cell responses assessed on day 21.

**(d)** Magnitude of PBMC CD8 T cell responses on day 21 as measured by intracellular cytokine staining for IFN-γ following WT Adpgk peptide stimulation *in vitro* for 6 hours (left panel) and tetramer staining with ASMTNMELM:H-2D$^b$-PE (right panel). Error bars are geometric mean with 95% CI.

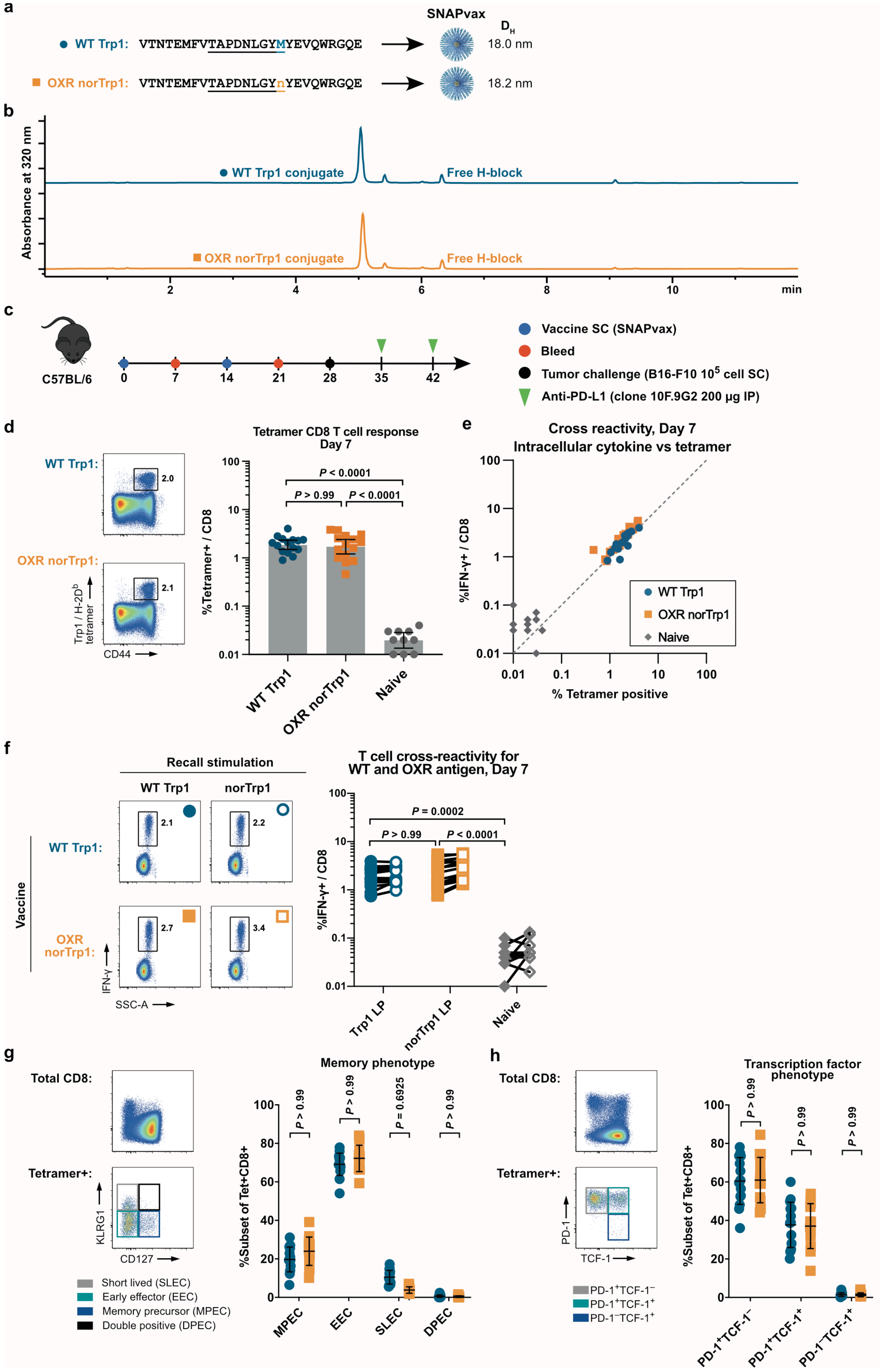
a
SNAPvax
WT Trp1: VTNTEMFVTAPDNLGYMYEVQWRGQE
OXR norTrp1: VTNTEMFVTAPDNLGYnYEVQWRGQE
18.0 nm
18.2 nm
b
Absorbance at 320 nm
WT Trp1 conjugate
Free H-block
OXR norTrp1 conjugate
min
c
C57BL/6
Vaccine SC (SNAPvax)
Bleed
Tumor challenge (B16-F10 10^5 cell SC)
Anti-PD-L1 (clone 10F.9G2 200 μg IP)
d
Tetramer CD8 T cell response
Day 7
%Tetramer+ / CD8
CD44
WT Trp1
OXR norTrp1
Naive
e
Cross reactivity, Day 7
Intracellular cytokine vs tetramer
%IFN-γ+ / CD8
% Tetramer positive
f
Recall stimulation
Vaccine
T cell cross-reactivity for
WT and OXR antigen, Day 7
%IFN-γ+ / CD8
SSC-A
Trp1 LP
norTrp1 LP
g
Total CD8:
Tetramer+:
Memory phenotype
%Subset of Tet+CD8+
KLRG1
CD127
Short lived (SLEC)
Early effector (EEC)
Memory precursor (MPEC)
Double positive (DPEC)
MPEC
EEC
SLEC
DPEC
h
Transcription factor
phenotype
PD-1
TCF-1

**Supplementary figure 5 | Norleucine substitution of methionine in Trp1 is equivalent to native Trp1**
(**a**) Sequence of tumor self-antigen Trp1 and norleucine-substituted Trp1 (OXR norTrp1). The 9-mer epitope known to bind MHC-I allele H-2D$^b$ is underlined and bold. The Methionine in Trp1 (blue M) is substituted with norleucine in norTrp1 (orange n) and is known to function as an MHC anchor residue. Each peptide was formulated as SNAPvax and formed uniform nanoparticles with the hydrodynamic diameter ($D_H$) shown.
(**b**) Reverse phase HPLC traces of SNAPvax conjugates of WT Trp1 (blue) and OXR norTrp1 (orange) with y-axis showing absorbance at 320 nm in arbitrary absorbance units (mAU). The two conjugates elute at similar times. As designed, an approximately 10% molar excess of free H-block is present in the formulation, which incorporates in the interior of the self-assembling nanoparticle micelle upon reconstitution in aqueous buffer.
(**c**) C57BL/6 mice (n = 10–15 per group) were vaccinated subcutaneously (SC) on day 0 and 14, bled on day 7 and 21, challenged subcutaneously with $10^5$ cells of B16-F10 on day 28, and treated with anti-PD-L1 on day 35 and 42. Data presented in (d-h).
(**d**) CD8 T cell responses on day 7 as measured by tetramer staining with TAPDNLGYM:H-2D$^b$-PE. Histograms show concatenated samples of all acquired events for each vaccine group (left), and bar graph shows quantification of each individual sample (right). *P*-values assessed by one-way ANOVA (Kruskal-Wallis with Dunn's correction). Error bars are geometric mean with 95% CI.
(**e**) Whole blood from each mouse was split and either stimulated with WT peptide followed by intracellular cytokine staining to determine IFN-γ production or stained with WT Trp1-tetramer.
(**f**) CD8 T cells from whole blood were stimulated with either WT Trp1 (solid) or OXR norTrp1 (open) peptides**.** Histograms show concatenated samples of all acquired events for each condition (left), and individual samples quantified with lines connecting paired samples (right). P-values assessed by one-way ANOVA (Kruskal-Wallis with Dunn's correction).
(**g**) Memory phenotype of tetramer+ cells on day 7. Histograms show concatenated samples of all acquired events for total CD8 T cells (top left) and tetramer+ CD8 T cells (bottom left), and bar graph shows quantification of each individual sample (right). P-values are by two-way ANOVA (Sidak's multiple comparison test) comparing responses between groups. Error bars are mean ± standard deviation.
(**h**) Transcription factor (TCF1) and activation status (PD-1) of tetramer+ cells on day 7. Histograms show concatenated samples of all acquired events for total CD8 T cells (top left) and tetramer+ CD8 T cells (bottom left), and bar graph shows quantification of each individual sample (right). P-values are by two-way ANOVA (Sidak's multiple comparison test) comparing responses between groups. Error bars are mean ± standard deviation.

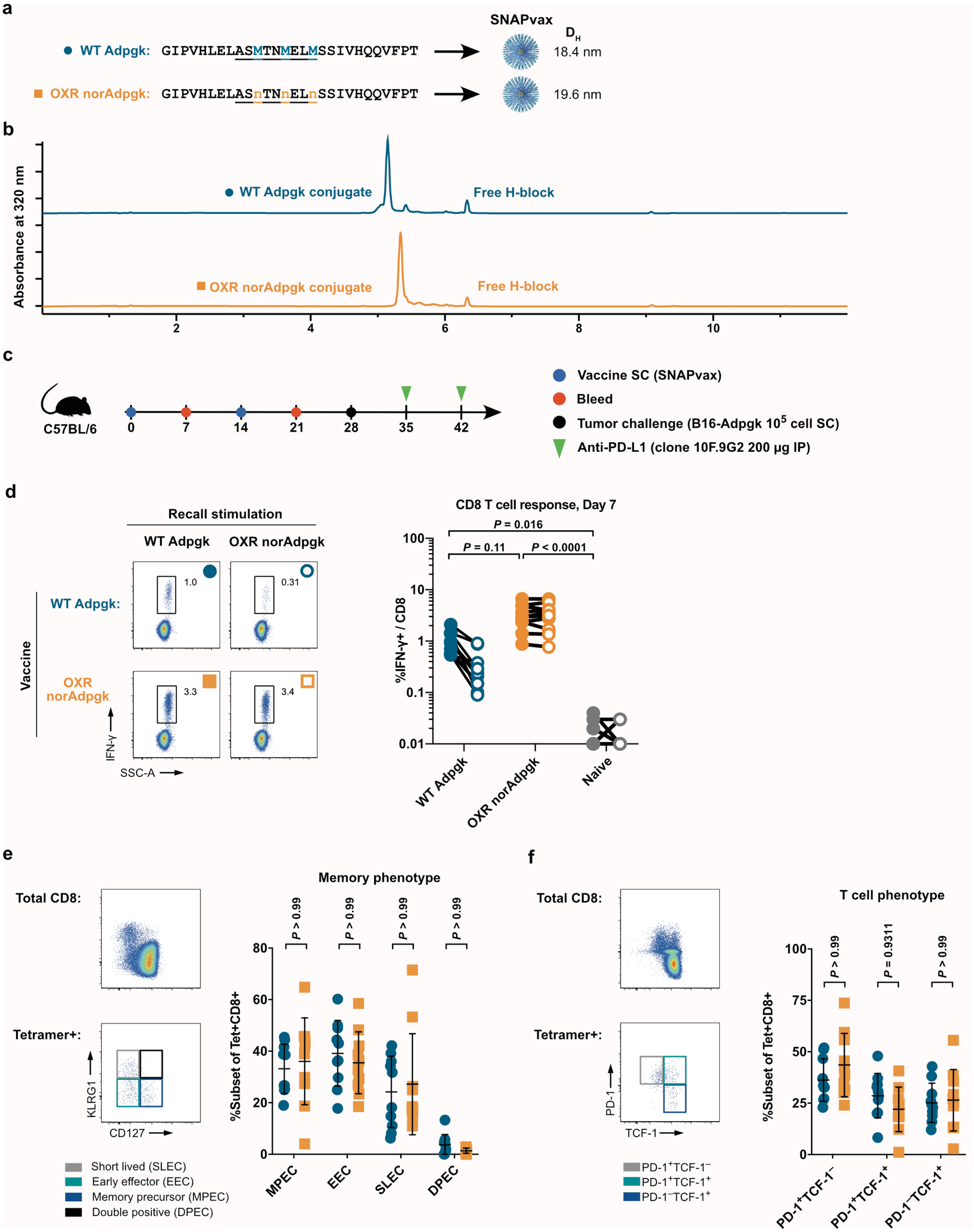
a
SNAPvax
D_H
WT Adpgk: GIPVHLELASMTNMELMSSIVHQQVFPT
18.4 nm
OXR norAdpgk: GIPVHLELASnTNnELnSSIVHQQVFPT
19.6 nm
b
Absorbance at 320 nm
WT Adpgk conjugate
Free H-block
OXR norAdpgk conjugate
Free H-block
2
4
6
8
10
c
C57BL/6
0
7
14
21
28
35
42
Vaccine SC (SNAPvax)
Bleed
Tumor challenge (B16-Adpgk 10^5 cell SC)
Anti-PD-L1 (clone 10F.9G2 200 µg IP)
d
Recall stimulation
WT Adpgk
OXR norAdpgk
Vaccine
WT Adpgk:
OXR norAdpgk
1.0
0.31
3.3
3.4
IFN-γ
SSC-A
CD8 T cell response, Day 7
P = 0.016
P = 0.11
P < 0.0001
%IFN-γ+ / CD8
100
10
1
0.1
0.01
WT Adpgk
OXR norAdpgk
Naive
e
Total CD8:
Tetramer+:
KLRG1
CD127
Short lived (SLEC)
Early effector (EEC)
Memory precursor (MPEC)
Double positive (DPEC)
Memory phenotype
P > 0.99
P > 0.99
P > 0.99
P > 0.99
%Subset of Tet+CD8+
80
60
40
20
0
MPEC
EEC
SLEC
DPEC
f
Total CD8:
Tetramer+:
PD-1
TCF-1
PD-1+TCF-1−
PD-1+TCF-1+
PD-1−TCF-1+
T cell phenotype
P > 0.99
P = 0.9311
P > 0.99
%Subset of Tet+CD8+
100
75
50
25
0
PD-1+TCF-1−
PD-1+TCF-1+
PD-1−TCF-1+

**Supplementary figure 6 | Norleucine substituted Adpgk is equivalent to native Adpgk**
(**a**) Sequence WT Adpgk and and norleucine-substituted Adpgk (OXR norAdpgk). The 9-mer epitope known to bind MHC-I allele H-2Db is underlined and bold. The three Methionine residues in Adpgk (blue M) are substituted with norleucine in norAdpgk (orange n). Each peptide was formulated as SNAPvax and formed uniform nanoparticles with the hydrodynamic diameter ($D_H$) shown.
(**b**) Reverse phase HPLC traces of SNAPvax conjugates of WT Adpgk (blue) and OXR norAdpgk (orange) with y-axis showing absorbance at 320 nm in arbitrary absorbance units (mAU). The two conjugates elute at similar times. As designed, an approximately 10% molar excess of free H-block is present in the formulation, which incorporates in the interior of the self-assembling nanoparticle micelle upon reconstitution in aqueous buffer.
(**c**) C57BL/6 mice (n = 10–15 per group) were vaccinated subcutaneously (SC) on day 0 and 14, bled on day 7 and 21, challenged subcutaneously with $10^5$ cells of B16-Adpgk on day 28, and treated with anti-PD-L1 on day 35 and 42.
(**d**) CD8 T cells from whole blood were stimulated with either WT Adpgk (solid) or OXR norAdpgk (open) peptides and assessed by intracellular cytokine staining for IFN-γ. *P*-values assessed by one-way ANOVA (Kruskal-Wallis with Dunn's correction).
(**e**) Memory phenotype of tetramer+ cells on day 7. Histograms show concatenated samples of all acquired events for total CD8 T cells (top left) and tetramer+ CD8 T cells (bottom left), and bar graph shows quantification of each individual sample (right). P-values are by two-way ANOVA (Sidak's multiple comparison test) comparing responses between groups. Error bars are mean ± standard deviation.
(**f**) Transcription factor (TCF1) and activation status (PD-1) of tetramer+ cells on day 7. Histograms show concatenated samples of all acquired events for total CD8 T cells (top left) and tetramer+ CD8 T cells (bottom left), and bar graph shows quantification of each individual sample (right). P-values are by two-way ANOVA (Sidak's multiple comparison test) comparing responses between groups. Error bars are mean ± standard deviation.

Abbreviations: MHC = major histocompatibility complex. WT = wildtype. OXR = oxidation resistant. nor = norleucine. SLEC = short lived effector. EEC = early effector. MPEC = memory precursor. DPEC = double positive. $D_h$ = hydrodynamic diameter.

**Supplementary Table 1: Peptide antigens**

| Name | Sequence | Protein | Cell line/virus |
|---|---|---|---|
| WT gp33 min | KAVYNFATC | glycoprotein 33 | LCMV |
| gp33 abut min | KAVYNFATB | glycoprotein 33 | LCMV |
| gp33 WT LP | VITGIKAVYNFATCGIFAL | glycoprotein 33 | LCMV |
| gp33 abut LP | VITGIKAVYNFATBGIFAL | glycoprotein 33 | LCMV |
| gp33 serine min | VITGIKAVYNFATSGIFAL | glycoprotein 33 | LCMV |
| gp33 serine LP | KAVYNFATS | glycoprotein 33 | LCMV |
| M27 WT min | GVELCPGNKYEM | Obscurin-like protein 1 | B16/F10 |
| M27 abut min | GVELBPGNKYEM | Obscurin-like protein 1 | B16/F10 |
| M27 WT LP | REGVELCPGNKYEMRRHGTTHSLVIHD | Obscurin-like protein 1 | B16/F10 |
| M27 abut LP | REGVELBPGNKYEMRRHGTTHSLVIHD | Obscurin-like protein 1 | B16/F10 |
| Trp1 WT min | TAPDNLGYM | Tyrosinase related protein 1 | B16/F10 |
| Trp1 nor min | TAPDNLGYn | Tyrosinase related protein 1 | B16/F10 |
| Trp1WT LP | VTNTEMFVTAPDNLGYMYEVQWPGQE | Tyrosinase related protein 1 | B16/F10 |
| Trp1 nor LP | VTNTEMFVTAPDNLGYnYEVQWPGQE | Tyrosinase related protein 1 | B16/F10 |
| Adpgk WT min | ASMTNMELM | ADP-dependent glucokinase | MC38 |

| | | | |
|---|---|---|---|
| Adpgk nor min | ASnTNnELn | ADP-dependent glucokinase | MC38 |
| Adpgk WT LP | GIPVHLELASMTNMELMSSIVHQQVFPT | ADP-dependent glucokinase | MC38 |
| Adpgk nor LP | GIPVHLELASnTNnELnSSIVHQQVFPT | ADP-dependent glucokinase | MC38 |
| Adpgk norvaline LP | GIPVHLELASvTNvELnSSIvHQQVFPT | ADP-dependent glucokinase | MC38 |
| Adpgk leucine LP | GIPVHLELASLTNLELLSSIVHQQVFPT | ADP-dependent glucokinase | MC38 |

B denotes alpha-aminobutyric acid; n denotes norleucine, v denotes norvaline

**Supplementary Table 2: Intracellular cytokine staining panel**

| Target | Fluorophore | Clone | Supplier | Cat # | Dilution | Step |
|---|---|---|---|---|---|---|
| LiveDead | UV Blue | NHS dye | ThermoFisher | L23105 | 800 | Pre-stain |
| CD4 | BUV395 | RM4-4 | BD | 740209 | 800 | Surface |
| CD8a | BUV805 | 53-6.7 | BD | 612898 | 400 | Surface |
| PD-1 | BV421 | EH12.2H7 | BioLegend | 329920 | 200 | Surface |
| KLRG1 | BV785 | 2F1/KLRG1 | BD | 138429 | 200 | Surface |
| CD127 | Cy5PE | A7R34 | BioLegend | 135016 | 100 | Surface |
| CD3e | Ax700 | 17A2 | BD | 751418 | 1000 | Intracellular |
| IFN-γ | APC | XMG1.2 | BioLegend | 505810 | 200 | Intracellular |
| IL-2 | PE | JES6-5H4 | BD | 554428 | 400 | Intracellular |
| TNF-a | BV650 | MAb11 | BD | 563418 | 800 | Intracellular |

**Supplementary Table 3: Tetramer staining panel**

| Target | Fluorophore | Clone | Supplier | Cat # | Dilution | Step |
|---|---|---|---|---|---|---|
| LiveDead | UV Blue | NHS dye | ThermoFisher | L23105 | 800 | Pre-stain |
| Tetramer* | SAV-PE | -- | Invitrogen | SNN1007 | * | Surface |
| CD4 | BUV395 | RM4-4 | BD | 740209 | 800 | Surface |
| CD8a | APC-eF780 | 53-6.7 | ThermoFisher | 47-0081-82 | 400 | Surface |
| CD44 | BUV737 | IM7 | BD | 564392 | 1000 | Surface |
| PD-1 | BV421 | EH12.2H7 | BioLegend | 329920 | 200 | Surface |
| KLRG1 | BV785 | 2F1/KLRG1 | BD | 138429 | 200 | Surface |
| CD127 | Cy5PE | A7R34 | BioLegend | 135016 | 100 | Surface |
| CD3e | Ax700 | 17A2 | BD | 751418 | 1000 | Intracellular |
| Tcf1 | Ax647 | C63D9 | Cell Signaling Technology | 6709S | 1500 | Intracellular |

*Individual biotinylated pMHC unimers were formed into tetramers by mixing with streptavidin (SAV)-PE immediately prior to staining cells at a pMHC concentration of 0.1 µM

### METHODS

Methods and any associated references are available in the online version of the paper.

### NOTES

This work was supported in part by the Intramural Research Program of the VRC, NIAID, NIH and by Barinthus Biotherapeutics, North America. The findings and conclusions in this report are those of the authors and do not necessarily reflect the views of the funding agency, company or collaborators.

### ACKNOWLEDGEMENTS

The authors acknowledge the contributions of: Carmelo Chiedi, Sienna Rush, Oscar Hernandez, Gloria Salbador, and Diana Scorpio of the VRC Translational Research Program, NIH; Gloria Chu, Danielle Thompson, and Patricia Darrah of the Cellular Immunology Section, VRC, NIH.

### AUTHOR CONTRIBUTIONS

A.S.I. and G.M.L. conceived of the project. A.S.I. designed the experiments. A.S.I., A.M.S., M.M., M.E., and N.G.P performed the experiments. A.S.I., C.M.G., D.R.W. wrote the manuscript; R.A.S. and G.M.L. revised the manuscript. All authors had complete access to all data, contributed to the analysis, and reviewed the manuscript.

### COMPETING FINANCIAL INTERESTS

The authors declare competing financial interests: details are available in the online version of the paper.

A.S.I., C.M.G., A.R., Q.R., F.B., J.P.F., N.B., R.A.S., D.R.W., and G.M.L. are listed as inventors on patents describing peptide-based vaccines including those with OXR antigens. A.S.I., C.M.G., D.R.W., N.G.P., and G.M.L. are current or prior employees of Barinthus Biotherapeutics North America, Vaccitech North America or Avidea Technologies, Inc. which are commercializing peptide-based drug delivery technologies for immunotherapeutic applications.
